# Content-Aware Image Restoration: Pushing the Limits of Fluorescence Microscopy

**DOI:** 10.1101/236463

**Authors:** Martin Weigert, Uwe Schmidt, Tobias Boothe, Andreas Müller, Alexandr Dibrov, Akanksha Jain, Benjamin Wilhelm, Deborah Schmidt, Coleman Broaddus, Siân Culley, Mauricio Rocha-Martins, Fabián Segovia-Miranda, Caren Norden, Ricardo Henriques, Marino Zerial, Michele Solimena, Jochen Rink, Pavel Tomancak, Loic Royer, Florian Jug, Eugene W. Myers

## Abstract

Fluorescence microscopy is a key driver of discoveries in the life-sciences, with observable phenomena being limited by the optics of the microscope, the chemistry of the fluorophores, and the maximum photon exposure tolerated by the sample. These limits necessitate trade-offs between imaging speed, spatial resolution, light exposure, and imaging depth. In this work we show how image restoration based on deep learning extends the range of biological phenomena observable by microscopy. On seven concrete examples we demonstrate how microscopy images can be restored even if 60-fold fewer photons are used during acquisition, how near isotropic resolution can be achieved with up to 10-fold under-sampling along the axial direction, and how tubular and granular structures smaller than the diffraction limit can be resolved at 20-times higher frame-rates compared to state-of-the-art methods. All developed image restoration methods are freely available as open source software in Python, Fiji, and Knime.

## 1 Introduction

Fluorescence microscopy is an indispensable tool in the life sciences for investigating the spatio-temporal dynamics of cells, tissues, and developing organisms. Recent advances, such as light-sheet microscopy [1–3], structured illumination microscopy [4, 5], and super-resolution microscopy [6–8] enable time resolved volumetric imaging of biological processes within cells at high resolution. The quality at which these processes can be faithfully recorded, however, is not only determined by the spatial resolution of the used optical device, but also by the desired temporal resolution, the total duration of an experiment, the required imaging depth, the achievable fluorophore density, bleaching, and photo-toxicity [9, 10]. These aspects cannot all be optimized at the same time – one must make trade-offs, for example, sacrificing signal-to-noise ratio by reducing exposure time in order to gain imaging speed. Such trade-offs are often depicted by a *design-space* that has resolution, speed, light-exposure, and imaging depth as its dimensions (Figure 1a) with the volume being limited by the maximal photon budget compatible with sample health [11, 12].

**Figure 1:**
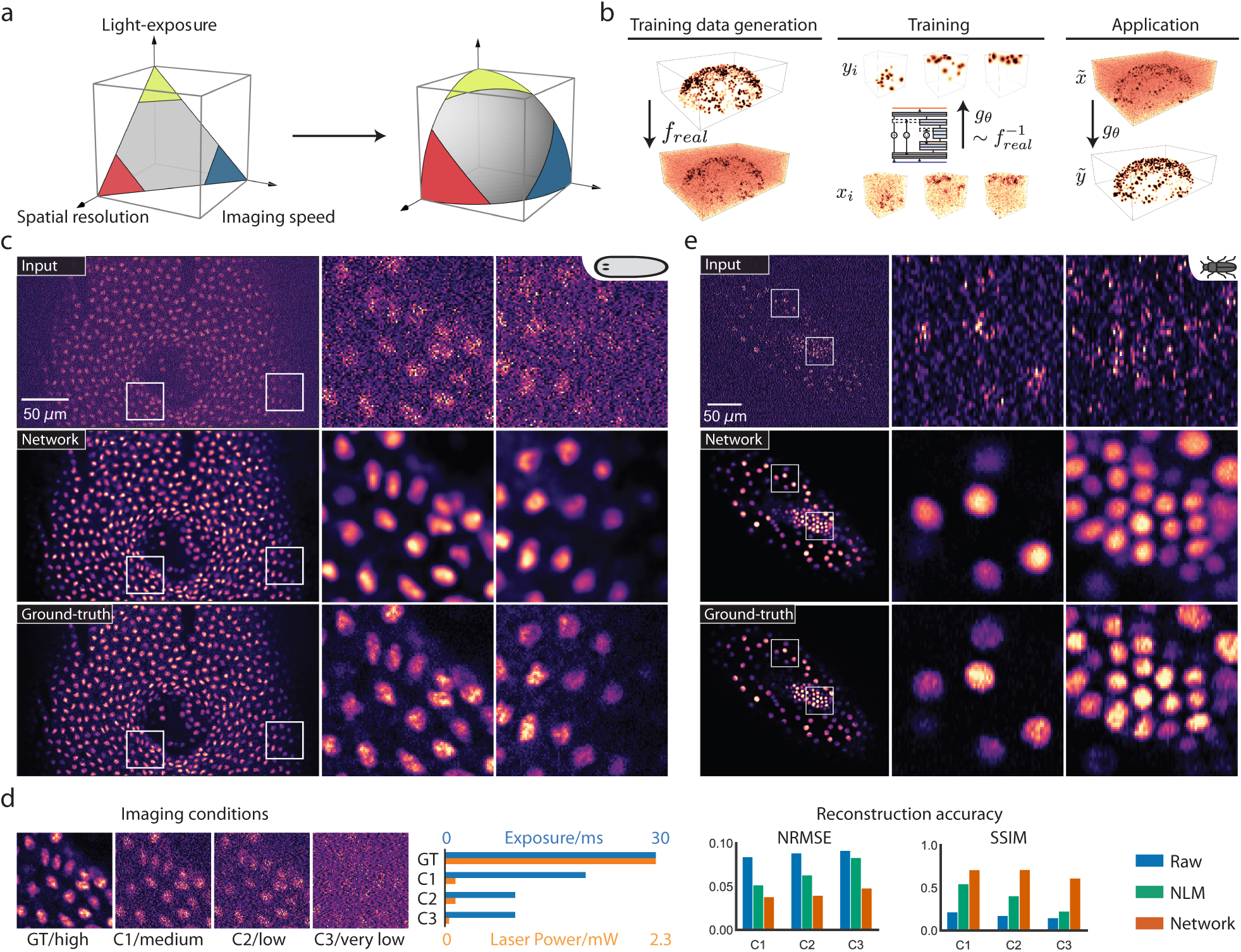
Content-aware image restoration in fluorescence microscopy. **(a)** The design-space of fluorescence microscopes. Trade-offs between imaging speed, spatial resolution, and light exposure need to be found to best capture a given sample within the constraints of a maximal photon budget (we omit imaging depth as the fourth dimension for illustrative purposes). Content-aware restoration (CARE) networks enlarge the design-space by restoring image aspects that suffered due to the trade-off used during imaging. **(b)** Restoration of noisy (low SNR) volumes. Pairs of high SNR and low SNR volumes are acquired at the microscope. Each pair (*x_i_, y_i_*) consists of two registered low and high SNR images of the same biological sample. A deep convolutional neural network is then trained to restore *y_i_* from *x_i_*. The trained CARE network is then applied to previously unseen, potentially very low SNR images *x*˜, yielding restored images *y*˜. **(c)** Input data and restorations for nucleus-stained (RedDot1) flatworm (*Schmidtea mediterranea*). Shown are a single image-plane of a raw input stack (top row), the network prediction (middle row), and the high SNR gold-standard/ground-truth (bottom row). Due to the photo-sensitivity of the flatworm, ground-truth data can only be obtained from fixed samples. Once trained, CARE networks enable live-cell imaging of *Schmidtea mediterranea* for the first time. For a comparison to a total of eight denoising methods, please see Supp. Figure 4. Details on training data and network parameters are given in Supp. Table 3. **(d)** Quantification of prediction error for *Schmidtea mediterranea* for different laser intensities and exposure times (C1 to C3). Bar plots show normalized root-mean-squared error (NRMSE) and structural similarity (SSIM, higher is better) for the input, for a denoising baseline (NLM [22]), and for our content-aware restorations. **(e)** Input data and restorations for a nucleus-labeled (EFA::nGFP) red flour beetle (*Tribolium castaneum*) embryo, again showing a single image-plane of the raw input data (top row), the network prediction (middle row), and the high SNR ground-truth data (bottom row). Please see Supp. Table 3 for training and network details.

These trade-offs can be addressed by optimizing the microscopy hardware, yet there are physical limits that cannot easily be overcome. Therefore, computational procedures to improve the quality of acquired microscopy images are becoming increasingly important. For instance, in the above-mentioned trade-off between exposure and speed, one could apply computational image restoration to maintain an image quality that is still sufficient for downstream data quantification at high acquisition speed. Super resolution microscopy [4, 13–16], deconvolution [17–19], surface projection algorithms [20, 21], and denoising methods [22–24] are examples of sophisticated image restoration algorithms that can push the limit of the design-space, and thus allow one to recover important biological information that would be inaccessible by imaging alone. Most common image restoration problems, however, have multiple possible solutions, and require additional assumptions in order to select one solution as the final restoration. These assumptions are typically general, *e.g*. requiring certain level of smoothness of the restored image, and therefore are not dependent on the specific content of the images to be restored. Intuitively, a method that leverages available knowledge about the data at hand ought to reach superior restoration results.

Deep Learning (DL) is such a method, since it can learn to perform complex tasks on specific data [25, 26]. It employs large multi-layered neural networks that compute results after being trained on annotated example data (*i.e*. gold-standard, *ground-truth* data). Spectacular results reaching human-level performance have for example been achieved on the classification of natural images [27]. In biology, DL methods have for instance been applied to the automatic extraction of connectomes from large electron microscopy data [28], for classification of image-based high-content screens [29], fluorescence signal prediction from label-free images [30, 31], resolution enhancement in histopathology [32], or for single molecule localization in super resolution microscopy [33, 34]. However, the direct application of DL methods to image restoration tasks in fluorescence microscopy is complicated by the absence of training data sets and the impossibility of generating them manually.

In this paper, we present a solution to the problem of missing training data for DL in fluorescence microscopy by developing strategies to generate such data. This enables us to apply neural networks to image restoration tasks, such as image denoising, surface projection, recovery of isotropic resolution, and the restoration of sub-diffraction structures. We show, in a variety of imaging scenarios, that trained *content-aware restoration* (CARE) networks produce results that were previously unobtainable. This means that the application of CARE to biological images allows to transcend the limitations of the design-space (Figure 1a), pushing the limits of the possible in fluorescence microscopy through machine learned image computation.

## 2 Results

In fluorescence microscopy one is often forced to image samples at low signal intensities, resulting in difficult to analyze, low signal-to-noise ratio (SNR) images. One way to improve SNR is to increase laser power or exposure times which, unfortunately, is usually detrimental to the sample, limiting the possible duration of the recording and introducing artifacts due to photo-damage. An alternative solution is to image at low SNR, and later computationally restore acquired images. Classical approaches, such as Non-local-means denoising [22], can in principle achieve this, but without leveraging available knowledge about the data at hand.

To address this problem with machine learning, we developed *content-aware* image *re*storation (CARE) networks, adapted to a specific experimental setup, hypothesizing that they produce results superior to classical, content-agnostic methods. In the case of image denoising, we acquired pairs of images at low and high signal-to-noise ratios, used them as input and ground-truth to train CARE networks, and applied the trained networks to remove noise in previously unseen data.

### Image Restoration with Physically Acquired Training Data

To demonstrate the power of this approach in biology, we applied it to the imaging of the flatworm *Schmidtea mediterranea*, a model organism for studying tissue regeneration. This organism is exceptionally sensitive to even moderate amounts of laser light [35], suffering muscle flinching at desirable illumination levels even when anesthetized (Supp. Video 1). Using a laser power that reduces flinching to an acceptable level results in images with such low SNR that they are impossible to interpret directly. Consequently, live imaging of *S. mediterranea* has thus far been intractable.

To address this problem with CARE, we imaged *fixed* worm samples at several laser intensities. We acquired well-registered pairs of images, a low-SNR image at laser power compatible with live imaging, and a high-SNR image, serving as ground-truth. We then trained a convolutional neural network^1^ and applied the trained network to previously unseen live imaging data of *S. mediterranea*. We consistently obtained high quality restorations, even if the SNR of the images was very low, *e.g*. being acquired with a 60-fold reduced light-dosage (Figure 1c, Supp. Video 2, Supp. Figure 1–3). To quantify this observation, we measured the restoration error between prediction and ground-truth images for three different exposure and laser-power conditions. Both, the NRMSE^2^ and the SSIM^3^ measures of error improved considerably when compared to results obtained by several potent classical denoising methods (Figure 1d, Supp. Figure 2 & 4, Supp. Table 1). We further observed that already a small number of training images (*e.g*. 200 patches of size 64×64×16) leads to an acceptable image restoration quality (Supp. Figure 5). Moreover, while training a CARE network can take several hours, the restoration time for a volume of size 1024×1024×100 was less than 20 seconds on a single graphics processing unit^4^. In this case, CARE networks are able to take input data that are unusable for biological investigations and turn them into high-quality time-lapse data, providing the first practical framework for live-cell imaging of *S. mediterranea*.

We next asked whether CARE improves common downstream analysis tasks in live-cell imaging, such as nuclei segmentation. We used light-sheet recordings of developing *Tribolium castaneum* (red flour beetle) embryos, and as before trained a network on image pairs of samples acquired at high and low laser powers (Figure 1e). The resulting CARE network performs well even on extremely noisy, previously unseen live-imaging data, acquired with up to 70-fold reduced light-dosage compared to typical imaging protocols [39] (Supp. Chapter 4, Supp. Video 3, Supp. Figure 6). In order to test the benefits of CARE for segmentation, we applied a simple nuclei segmentation pipeline to raw and restored image stacks of *T. castaneum*. The results show that compared to manual expert segmentation, the segmentation accuracy (as measured with the standard SEG score [40]) improved from SEG = 0.47 on the classically denoised raw stacks to SEG = 0.65 on the CARE restored volumes (Supp. Figure 7). Since this segmentation performance is achieved at significantly reduced laser power, the gained photon budget can now be spent on the imaging speed and light-exposure dimensions of the design-space. This means that *Tribolium* embryos, when restored with CARE, can be imaged longer and at higher frame rates, enabling improved tracking of cell lineages.

Encouraged by the performance of CARE on two independent denoising tasks, we asked whether such networks can also solve more complex, composite tasks. In biology it is often useful to image a 3D volume and project it to a 2D surface for analysis, for example when studying cell behavior in developing epithelia of the fruit fly *Drosophila melanogaster* [41–43]. Also in this context, it is beneficial to optimize the trade-off between laser-power and imaging speed, usually resulting in rather low-SNR images. Thus, this restoration problem is composed of projection and denoising, presenting the opportunity to test if CARE networks can deal with such composite tasks. For training, we again acquired pairs of low and high SNR 3D image stacks, and further generated 2D projection images from the high SNR stacks [20] that serve as ground-truth (Figure 2a). We developed a task-specific network architecture that consists of two jointly trained parts: a network for surface projection, followed by a network for image denoising (Figure 2b, Supp. Figure 12 and Supp. Chapter 2). The results show that with CARE, reducing light dosage up to 10-fold has virtually no adverse effect on the quality of segmentation and tracking results obtained on the projected 2D images with an established analysis pipeline [44] (Figure. 2 c & d, Supp. Video 4, and Supp. Figure 9, 10 & 11). Even for this complex task, the gained photon-budget can be used to move beyond the design-space, for example by increasing temporal resolution, and consequently improving the precision of tracking of cell behaviors during wing morphogenesis [44].

**Figure 2:**
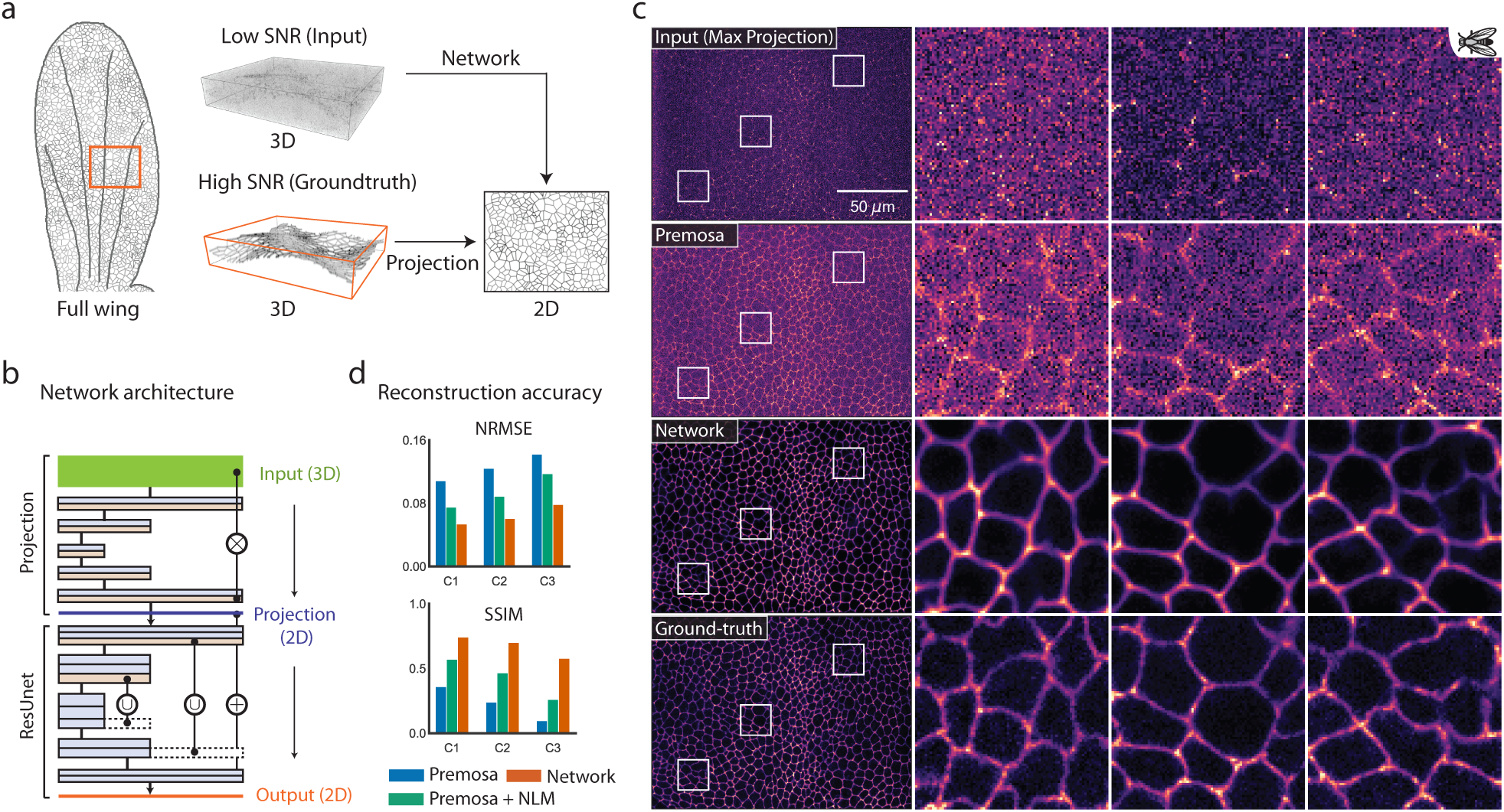
A CARE network jointly solving the composite task of surface projection and denoising. **(a)** Schematic of the composite task. A developing *Drosophila* wing is a single layer of cells, embedded in a 3D volume. Imaging of long time-lapses requires to image at low SNR to avoid photo-toxicity and bleaching. With CARE, the cell layer of interest can be projected onto a 2D image, while also being denoised by the same, composite network. **(b)** The architecture of the proposed CARE network consists of two consecutive sub-networks: the first one performing the projection of voxel intensities (top half), and the second one denoising this projection. **(c)** Results obtained by the proposed CARE network on E-cadherin labeled fly wing data. Shown is a max-projection of the raw input data (top row), result obtained by applying the state-of-the-art projection method Premosa [20] (second row), the solution computed by our trained network (third row), and the desired (ground-truth) projection obtained by applying Premosa on a very high laser-power (high SNR) acquisition of the same sample. Details on training data and network parameters can be found in Supp. Table 3. The ground-truth data was obtained at a laser intensity that cannot be used for live-cell imaging without causing damage to the sample (see main text for details). **(d)** Quantification of restoration errors for data acquired at different laser intensities and exposure times (conditions C1–C3). We show normalized root-mean-square error (NRMSE) and structural similarity (SSIM, higher is better) between ground-truth images and results obtained using Premosa (blue), Premosa with additional denoising (NLM [22], green), and our trained CARE network (orange). For a comparison to additional baseline methods, please see Supp. Figure 10.

### Image Restoration with Semi-synthetic Training Data

Thus far, the application of CARE has relied on the availability of matching pairs of high and low quality images, both physically acquired at a microscope. However, this kind of data is not always available. Therefore, we investigated whether image pairs useful for training can be obtained also by computationally modifying existing microscopy images.

A common problem in fluorescence microscopy is that the axial resolution of volumetric acquisitions is significantly lower than the lateral resolution^5^. This *anisotropy* compromises the ability to accurately measure properties such as the shapes or volumes of cells. Anisotropy is caused by the inherent axial elongation of the optical point spread function (PSF), and the often low axial sampling rate of volumetric acquisitions required for fast imaging. For the restoration of anisotropic image resolution, adequate pairs of training data cannot directly be acquired at the microscope. Rather, we took well-resolved lateral slices as ground truth, and computationally modified them (*i.e*. applied a realistic imaging model, Supp. Chapter 2) to resemble anisotropic axial slices of the same image stack. In this way, we generated matching pairs of images showing the same content at axial and lateral resolutions. These *semi-synthetically* generated pairs are suitable to train a CARE network that then restores previously unseen axial slices to nearly isotropic resolution (Figure 3a, Supp. Figure 19, Supp. Chapter 2, and [46, 47]). In order to restore entire anisotropic volumes, we applied the trained network to all lateral image slices, taken in two orthogonal directions, averaged to a single isotropic restoration (Supp. Chapter 2).

**Figure 3:**
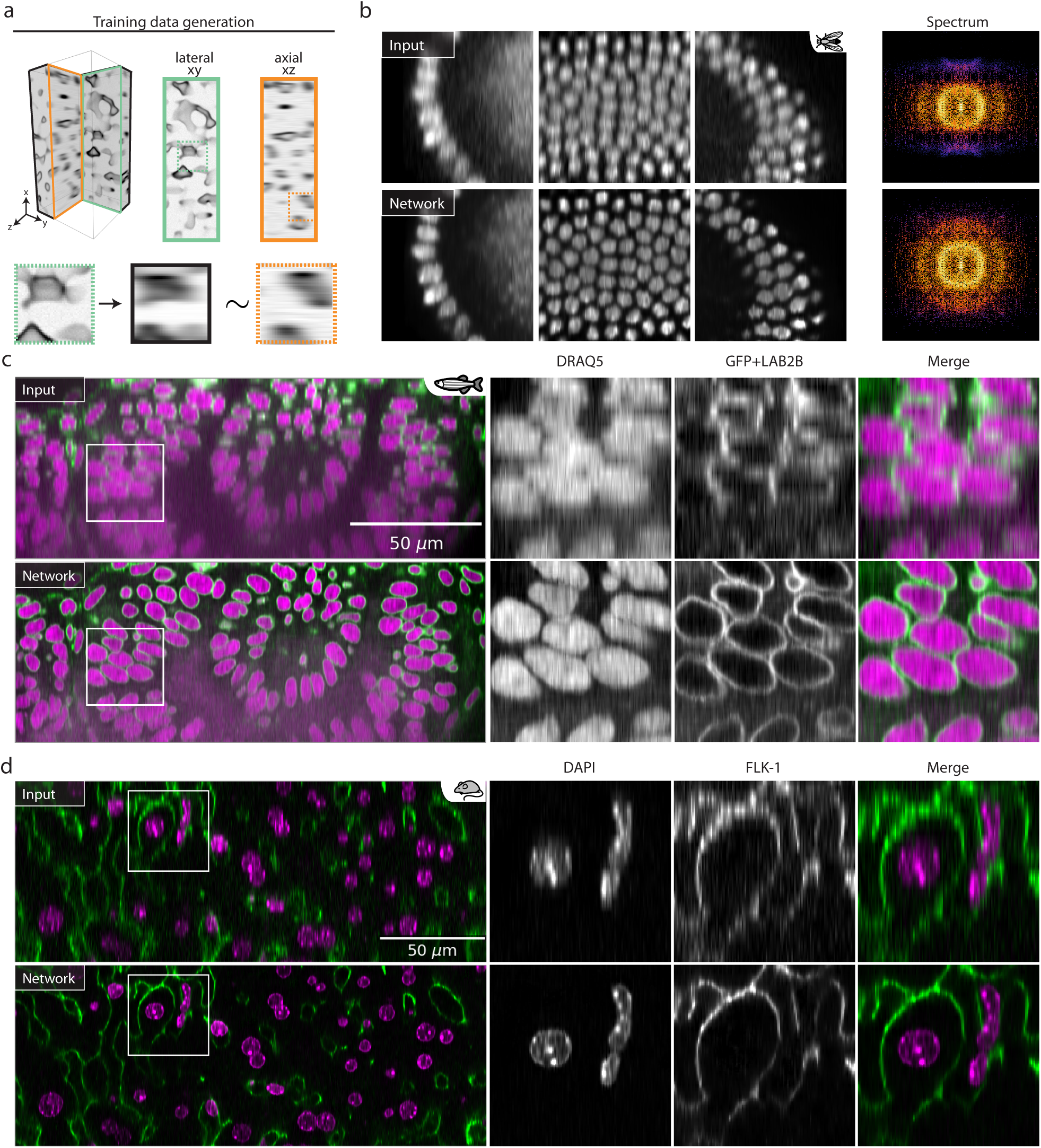
Isotropic restoration of 3D volumes with CARE. **(a)** Schematic of the semi-synthetic generation of training data. Lateral slices of the raw data (mint green) are used as ground-truth data. Corresponding anisotropic axial slices can be generated synthetically by down-sampling and convolution with the axial component of the PSF of the mimicked microscope (black inset). Raw axial slices (orange inset) cannot be used to train CARE networks because the isotropic image content is unknown. **(b)** Application of CARE on raw time-lapse acquisitions of *Drosophila melanogaster* [48]. Shown are three areas of raw axial input data (top row), and the respective isotropic restorations (bottom row). The rightmost column shows the Fourier-spectrum of raw and restored images and shows how missing parts of the spectrum are recovered by CARE. **(c)** An axial slice through a zebrafish retina in the anisotropic raw data (top row) and the isotropic restoration with CARE. Nuclei are DRAQ5 labeled in magenta and the nuclear envelope is labeled by GFP+LAB2B in green. For a comparison to deconvolution using Huygens (Scientific Volume Imaging, http://svi.nl), please see Supp. Figure 10. **(d)** An axial slice through mouse liver tissue, again showing the anisotropic raw data in the top row and the isotropic restoration below. Nuclei are labeled with DAPI and displayed in magenta, while sinusoids are labeled with FLK-1 and shown in green. Details on training data and network parameters for all cases can be found in Supp. Table 3.

We applied this strategy to increase axial resolution of acquired volumes of fruit fly embryos [48], zebrafish retina [49], and mouse liver, imaged with different fluorescence imaging techniques. The results show that CARE improved the axial resolution in all three cases considerably (Figure 3b-d, Supp. Video 5 & 6, and Supp. Figure 13 & 17). In order to quantify this, we performed Fourier-spectrum analysis of *Drosophila* volumes before and after restoration, and showed that the frequencies along the axial dimension are fully restored, while frequencies along the lateral dimensions remain unchanged (Supp. Figure 14). Since the purpose of the fruit fly data is to segment and track nuclei, we applied a common segmentation pipeline [50] to the raw and restored images, and observed that the fraction of incorrectly identified nuclei was lowered from 1.7% to 0.2% (Supp. Chapter 2, Supp. Figure 15 & 16). Thus, restoring anisotropic volumetric embryo images to effectively isotropic stacks, leads to improved segmentation, and will enable more reliable extraction of developmental lineages.

The zebrafish and mouse liver data are examples of live and fixed two-channel imaging of large organs, both requiring high imaging speed and isotropic resolution for downstream analysis. While isotropy facilitates segmentation and subsequent quantification of shapes and volumes of cells, vessels, or other biological objects of interest, higher imaging speed enables imaging of larger volumes and their tracking over time. Indeed, respective CARE networks deliver the desired axial resolution with up to 10-fold fewer axial slices (Figure 3 c & d, see Supp. Figure 18 for comparison with classical deconvolution), allowing one to reach comparable results ten times faster. Moreover, we observed that for these two-channel data sets, the network learned to exploit correlations between channels, leading to a better overall restoration quality compared to results based on individual channels (Supp. Figure 17).

Taken together, increasing isotropic resolution through CARE networks, trained on semi-synthetic pairs of images, benefits both imaging speed and accuracy of downstream analysis in many biological applications. Moreover, since training data can computationally be derived from the data to be restored, this method can be applied to any already acquired data set.

### Image Restoration with Synthetic Training Data

Having seen the potential of using semi-synthetic training data for CARE, we next investigated whether reasonable restorations can be achieved from synthetic image data alone, *i.e*. without involving real microscopy data during training.

In most of the previous applications, one of the main benefits of CARE networks was improved imaging speed. Many biological applications additionally require resolving sub-diffraction structures in the context of live-cell imaging. Super-resolution imaging modalities achieve the necessary resolution, but suffer from low acquisition rates. On the other hand, widefield imaging offers the necessary speed, but lacks the required resolution. We therefore tested whether CARE can computationally resolve sub-diffraction structures using only widefield images as input. Note that this is a fundamentally different approach compared to recently proposed methods for single molecule localization microscopy that reconstruct a single super-resolved image from multiple diffraction limited input frames using deep-learning [33, 34]. To this end, we developed *synthetic* generative models of tubular and point-like structures that are commonly studied in biology. In order to obtain synthetic image pairs, suitable for training CARE networks, we used these generated structures as ground-truth, and computationally modified them to resemble actual microscopy data (Supp. Chapter 2, Supp. Figure 21). Specifically, we created synthetic ground-truth images of tubular meshes resembling microtubules, and point-like structures of various sizes mimicking secretory granules. Then we computed synthetic input images by simulating the image degradation process by applying a PSF, camera noise, and background auto-fluorescence (Figure 4a, Supp. Chapter 2, and Supp. Figure 21). Finally, we trained a CARE network on these generated image pairs, and applied it to 2-channel widefield time-lapse images of rat INS-1 cells where the secretory granules and the microtubules were labeled (Figure 4b). We observed that the restoration of both microtubules and secretory granules exhibit a dramatically improved resolution, revealing structures imperceptible in the widefield images (Supp. Video 7, and Supp. Figure 20). To substantiate this observation, we compared the CARE restoration to the results obtained by deconvolution^6^, which is commonly used to enhance widefield images (Figure 4b). Line profiles through the data show the improved performance of CARE network over deconvolution (Figure 4b). We additionally compared results obtained by CARE with super-resolution radial fluctuations (SRRF [14]) results, a state-of-the-art method for reconstructing super-resolution images from widefield time-lapse data. We applied both methods on time-lapse widefield images of GFP-tagged microtubules in HeLa cells. The results show that both CARE and SRRF are able to resolve qualitatively similar microtubular structures (Figures 4c, Supp. Video 8). However, CARE reconstructions are at least 20 times faster, since they are computed from a single average of up to 10 consecutive raw images while SRRF required about 200 consecutive widefield frames.

**Figure 4:**
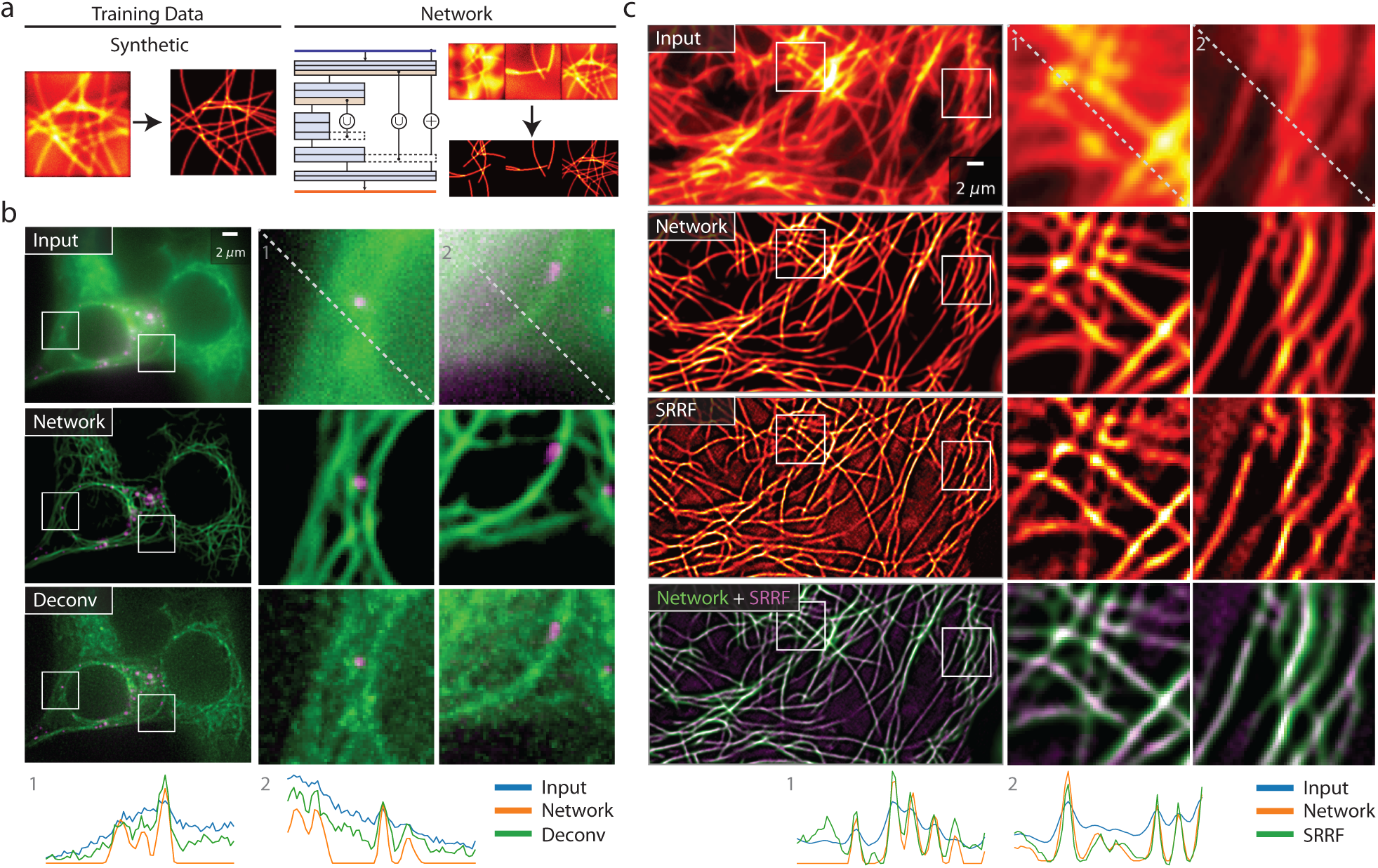
Resolving sub-diffraction structures in fluorescence microscopy images at very high frame rates with CARE. **(a)** Schematic of the fully-synthetic generation of training data pairs. Synthetic images of tubular and point-like structures where computationally generated as ground-truth data, and have further been processed to resemble actual microscopy data. CARE networks trained on such data can then be applied on real microscopy images of diffraction-limited tubular and point-like structures. **(b)** Raw widefield images of rat secretory granules (pEG-hIns-SNAP, magenta) and microtubules (SiR-tubulin, green) in insulin-secreting INS-1 cells (top row), the corresponding network restorations (second row), and a deconvolution result of the raw image as a baseline (bottom row). Below the images we show line-plots along the diagonal of both insets. **(c)** GFP-tagged microtubules in HeLa cells. Raw input image (top row), network restorations (second row), super-resolution images created by the state-of-the-art method SRRF [14] (third row), and a superposition of our results with restorations by SRRF (bottom row). Below the images we show line-plots along the diagonal of both insets.

Taken together, these results suggest that for structures that are straight-forward to model, such as microtubules, CARE networks can enhance widefield images to a resolution usually only obtainable with super-resolution microscopy, yet at considerably higher frame rates.

### Reliability of Image Restoration

We have shown that with the right training data, CARE networks perform remarkably well on a wide range of image restoration tasks, opening new avenues for biological observations. However, as for any image processing method, the issue of reliability of results needs to be addressed.

CARE networks are trained for a specific biological organism, fluorescent marker and microscope setting. When applying a network to data it was not trained for, results are likely to suffer in quality^7^. Nevertheless, we observed only minimal “hallucination” effects, where structures seen in the training data erroneously appear in restored images (see Supp. Figure 25). It was most pronounced when training and test data considerably differ in resolution while containing very specific structures that exhibit little variability (see Supp. Figure 26c). Otherwise, we observe similar effects only in very rare cases where structures are either so dim that they are not longer manifesting themselves in the input images, or in even rarer cases where the background noise can be interpreted as very low-SNR structure (see Supp. Figure 26a, showing the two strongest errors across the entire body of available image data). Naturally, it would be desirable to identify cases where the above-mentioned problems occur.

Therefore, to facilitate the evaluation of reliability of CARE network predictions, we changed the last network layer so that it predicts a *probability distribution*^8^ for each pixel (Figure 5a and Supp. Chapter 3. This distinguishes CARE from conventional image restoration approaches such as deconvolution [17, 51], where only a single restored intensity value is computed per pixel. For probabilistic CARE networks, the mean of the distribution is used as the restored pixel value, while the width (variance) of each pixel distribution encodes the *uncertainty* of pixel predictions. Intuitively, narrow distributions signify high confidence, whereas broad distributions indicate low confidence pixel predictions. This allows us to provide per-pixel confidence intervals of the restored image (Figure 5a, and Supp. Figure 22 & 23). These confidence intervals carry information about the reliability of CARE network predictions. We observed that variances tend to increase with restored pixel intensities. This makes it hard to intuitively understand which areas of an restored image are reliable or unreliable from a static image of per-pixel variances. Therefore, we visualize the uncertainty in short video sequences, where pixel intensities are randomly sampled from their respective distributions (Supp. Video 9). To a human observer, strong flicker in such videos highlights the areas where the uncertainty of image restorations is high. In the context of machine learning the accuracy can often be increased by aggregating several trained predictors [52]. In addition, we reasoned that by analyzing the consistency of network predictions we can assess their reliability. To that end, we train *ensembles* (Figure 5b) of about 5 CARE networks on randomized sequences of the same training data. We introduced a measure 𝒟 that quantifies the probabilistic ensemble disagreement per pixel (Supp. Chapter 3). 𝒟 takes values between 0 and 1, with higher values signifying larger disagreement, *i.e*. smaller overlap among the distributions predicted by the networks in the ensemble. Using fly wing denoising as an example, we observed that in areas where different networks in an ensemble predicted very similar structures, the disagreement measure 𝒟 was low (Figure 5c, top row), whereas in areas where the same networks predicted obviously dissimilar solutions, the corresponding values of 𝒟 were large (Figure 5c, bottom row). Therefore, training ensembles of CARE networks is useful to detect problematic image areas that cannot reliably be restored^9^.

**Figure 5:**
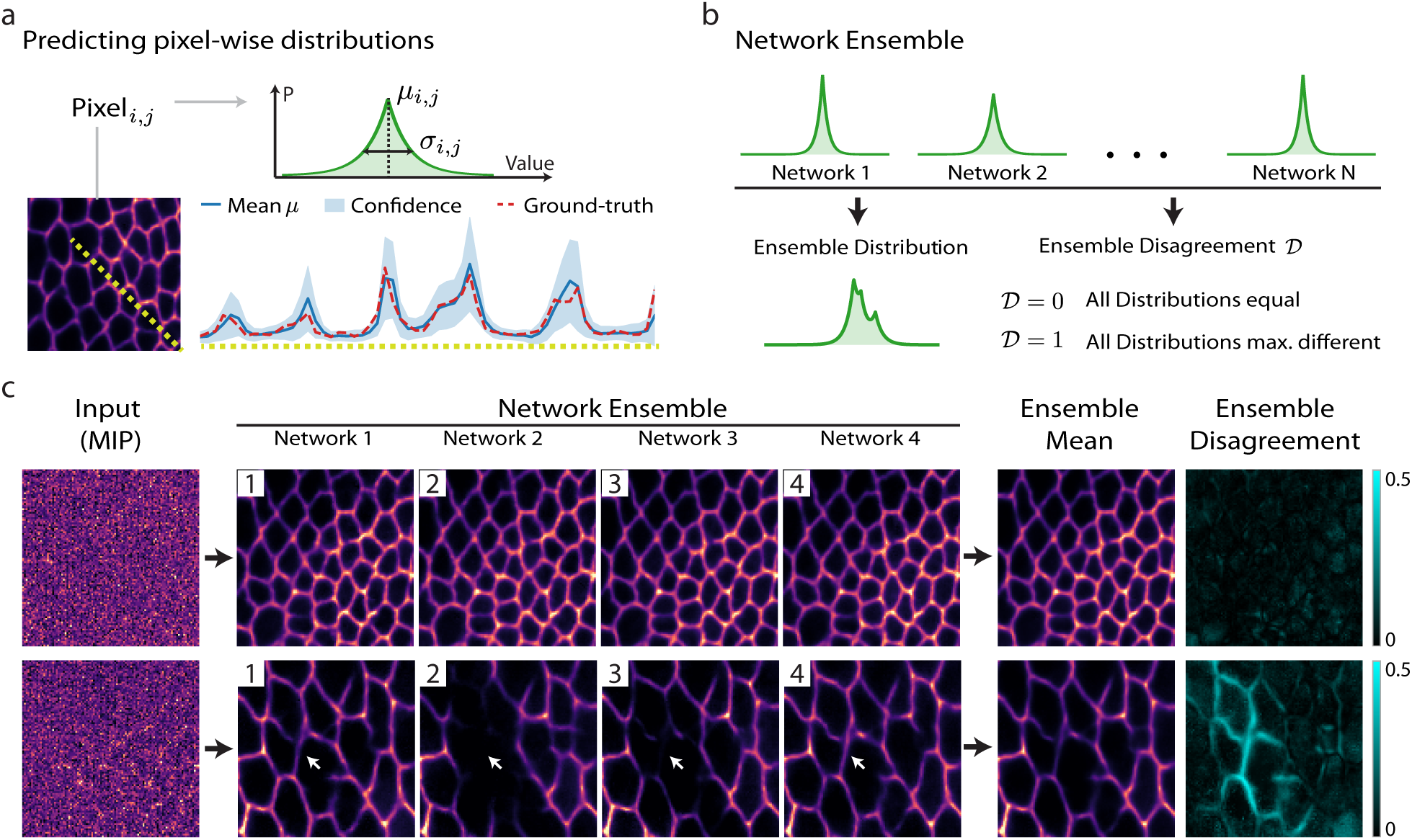
Quantifying the reliability of image restoration with CARE. **(a)** For every pixel of the restored image, CARE networks predict a (Laplace) distribution parameterized by its mean *μ* and variance or scale σ (top). These distributions provide pixel-wise confidence intervals (bottom), here shown for a surface projection and denoising network (*cf.* Figure 2). The line-plot shows predicted mean (blue) with 90% confidence interval (light-blue) and corresponding ground-truth (dashed red). **(b)** Multiple independently-trained CARE networks are combined to form an ensemble, resulting in an ensemble distribution and an ensemble disagreement measure𝒟 ∈ [0, 1]. **(c)** Ensemble predictions can vary, especially on challenging image regions. Shown are two examples for a surface projection and denoising CARE ensemble of 4 networks; from left to right: maximum projection of the input, the four individual network predictions, the ensemble mean, and the ensemble disagreement. The first example (top row) shows an image region with low disagreement, whereas the second example (bottom row) depicts a region where individual network predictions are substantially different, resulting in a high disagreement score in the affected image areas. For a detailed description of the theoretical foundation of above ideas, please refer to Supp. Chapter 3.

### Availability of Proposed Methods

Code for network training and prediction (written in Python using Keras [53] and TensorFlow [54]) is publicly available^10^. Furthermore, to make our restoration models readily available, we developed user-friendly FIJI plugins and KNIME workflows (Supp. Figure 27 & 28).

## 3 Discussion

We have introduced content-aware image restoration (CARE) networks designed to restore fluorescence microscopy data. A key feature of our approach is that generating training data does not require laborious manual training data generation. Application of CARE to raw images significantly expands the realm of observable biological phenomena. With CARE, flatworms can be imaged without unwanted muscle contractions, beetle embryos can be imaged much gentler and therefore longer and faster, large tiled scans of entire *Drosophila* wings can be imaged and simultaneously projected at dramatically increased temporal resolution, isotropic restorations of embryos and large organs can be computed from existing anisotropic data, and sub-diffraction structures can be restored from widefield systems at high frame rate. In all these examples, CARE allows one to invest the photon budget saved during imaging into improvement of acquisition parameters relevant for a given biological problem, such as speed of imaging, photo-toxicity, isotropy, or resolution.

Whether an experimentalist is willing to make the above mentioned investment, depends on her trust that a CARE network is accurately restoring the image. This is a valid concern, that applies to any image restoration approach. What sets CARE apart is the availability of additional readouts, *i.e*. per-pixel confidence intervals and ensemble disagreement scores. While strong disagreement indicates untrustworthy predictions, the converse is not necessarily true since all networks could simply make the same or similar mistakes. Still, the proposed disagreement score allows users to identify image regions where restorations might not be accurate.

We have shown multiple examples where image restoration with CARE networks positively impacts downstream image analysis, such as segmentation and tracking needed for extracting developmental lineages. Interestingly, in the case of *Tribolium*, CARE improved segmentation by efficient denoising, whereas in the case of *Drosophila*, the segmentation was improved by increasing the isotropy of volumetric acquisitions. These two benefits are not mutually exclusive and could very well be combined. In fact, we have shown on data from developing *Drosophila* wings, that composite tasks can jointly be trained. Future explorations of jointly training composite networks will further broaden the applicability of CARE to complex biological imaging problems.

Yet, CARE networks cannot be applied to all existing image restoration problems. For instance, the proposed isotropic restoration relies on the implicit assumption that structures of interest do appear in arbitrary orientations and that the PSF is constant throughout the image volume^11^. Additionally, CARE cannot be used if ground-truth can neither be physically acquired nor synthetically generated. The synthetic generation of training data could, in general, benefit from recent advances in computer vision, such as generative adversarial networks (GANs) [55]. For single molecule localization in super resolution microscopy, GANs are already successfully used [34]. Since our aim is to enable even novice users to train CARE networks on their own data, we decided to use the robust and easy to train U-net architecture [36]. Furthermore, the disagreement score we introduced could be used to identify instances where training and test data are incompatible, *i.e*. when a CARE network is applied on data that contains biological structures absent from the training set.

Overall, our results show that fluorescence microscopes can, in combination with content aware restorations, operate at higher frame-rates, shorter exposures, and lower light intensities, while reaching higher resolution, and thereby improving downstream analysis. The technology described here is readily accessible to the scientific community through the open source tools we provide. We predict that the current explosion of image data diversity and the ability of CARE networks to automatically adapt to various image contents, will make such learning approaches prevalent for biological image restoration and will open new windows into the inner workings of biological systems across scales.

## Acknowledgements

The authors want to thank Philipp Keller (Janelia) who provided *Drosophila* data. We thank Suzanne Eaton (MPI-CBG), Franz Gruber and Romina Piscitello for sharing the expertise in fly imaging and providing fly lines. We thank Anke So¨nmez for cell culture work. We thank Marija Matejcic (MPI-CBG) for generating and sharing the LAP2B transgenic line Tg(bactin:eGFP-LAP2B). We thank Benoit Lombardot from the Scientific Computing Facility (MPI-CBG). We thank the following Services and Facilities of the MPI-CBG for their support: Computer Department, Light Microscopy Facility (LMF) and Fish Facility. This work was supported by the German Federal Ministry of Research and Education (BMBF) under the codes 031L0102 (de.NBI) and 031L0044 (Sysbio II). M.S. was supported by the German Center for Diabetes Research (DZD e.V.). R.H. and S.C. was supported grants from the UK BBSRC (BB/M022374/1; BB/P027431/1; BB/R000697/1), UK MRC (MR/K015826/1) and Wellcome Trust (203276/Z/16/Z).

## Author Contributions

M.W. and L.R. initiated the research. M.W. and U.S. designed and implemented the training and validation methods. U.S., M.W., and F.J. designed and implemented the uncertainty readouts. T.B., A.M., A.D., S.C., F.S.M., R.H., M.R.M., and A.J. collected experimental data. A.D., C.B., and F.J. performed cell segmentation analysis. T.B. performed analysis on flatworm data. U.S. and M.W. designed and developed the Python package. F.J., B.W., and D.S. designed and developed the FIJI and KNIME integration. E.W.M supervised the project. F.J., M.W., P.T., L.R., U.S., & E.W.M wrote the manuscript, with input from all authors.

## Footnotes

1 We use networks of moderate size (10^6^ parameters) based on the U-net architecture [36, 37], together with a per-pixel similarity loss (*e.g*. absolute error) (*cf.* Supp. Figure 8, Supp. Chapter 2 and Supp. Table 3)

2 Normalized root-mean-square error.

3 Structural similarity index, measuring the perceived similarity between two images [38].

4 We used a Nvidia Titan X GPU for all presented experiments.

5 Some complex modalities allow for (close to) isotropic acquisitions, *e.g*. multi-view light-sheet microscopy [19, 45].

6 We used the on-board DeltaVision OMX deconvolution procedure.

7 As is the case for any (supervised) method based on machine learning.

8 We chose a Laplace distribution for simplicity and robustness, see Supp. Chapter 3.

9 Another example for the utility of ensemble disagreement can be found in Supp. Figure 24.

10 https://github.com/CSBDeep/CSBDeep

11 This assumption is only approximately true, becoming increasingly worse when imaging deeper into tissues.

